# The heart-liver axis controls platelet turnover by hepatic STAT3 phosphorylation and TPO regulation after acute myocardial infarction

**DOI:** 10.64898/2026.09.14.751386

**Authors:** Friedrich Reusswig, Benjamin Tajdar, Simone Gorressen, Jens W. Fischer, Margitta Elvers

## Abstract

**Background:** Platelets play a critical role in thrombo-inflammation following acute myocardial infarction (AMI).

**Objectives:** Their impact on hepatic thrombopoietin (TPO) regulation after AMI remains poorly understood to date.

**Methods:** Wildtype, thrombocytopenic or GPVI deficient mice underwent ischemia/reperfusion (I/R) injury. Platelet activation and turnover, and hepatic expression of different receptors involved in TPO regulation were analyzed.

**Results:** Here, we identify a heart-liver axis that dynamically regulates platelet production post AMI. An elevated platelet turnover with increased reticulated as well as desialylated platelets early after AMI was detected. This was accompanied by the upregulation of specific receptors such as Asgr1/2 and IL-6R and increased phosphorylation of STAT3 in the liver and elevated numbers of megakaryocytes in spleen tissue. Consequently, increased TPO plasma levels at 24h post AMI were detected and platelet counts were rapidly restored after AMI. Platelet depletion induces a compensatory increase in hepatic TPO expression and plasma TPO levels, accompanied by dysregulated STAT3 signaling. This response contrasts with GPVI-deficient (*Gp6*^*-/-*^) mice, which exhibit no major alterations in TPO regulation, suggesting distinct mechanisms between acute thrombocytopenia and chronic platelet receptor deficiency in platelet homeostasis after AMI. These findings provide the first evidence of a direct heart-liver axis regulating platelet homeostasis after AMI, driven by inflammatory and hepatic signaling pathways.

**Conclusions:** Our study highlights a previously unrecognized role of the heart-liver axis in controlling platelet turnover, with hepatic STAT3 signaling as a key mediator. These insights enhance our understanding of post AMI platelet homeostasis and may be critical for future therapeutic strategies.

## 1 INTRODUCTION

The maintenance of a consistent platelet count is crucial within mammalian physiology to prevent bleeding complications and mitigate the risk of occlusive thrombus formation, which can lead to ischemic events [1]. According to recent data from the World Health Organization (WHO, 2024, August 7), ischemic heart diseases persist as a predominant global cause of mortality. Myocardial infarction (MI) is the most common ischemic event, where the recommended therapeutic approach involves immediate recanalization via primary percutaneous coronary intervention of the occluded vessel and the implementation of dual antiplatelet therapy (DAPT) comprising aspirin and P2Y12 inhibitors [2, 3]. In the initial post-myocardial infarction phase, an augmented proliferation of megakaryocytes and heightened platelet production occur, leading to an elevation of platelet counts [4], concomitant with intensified platelet degranulation [5]. Under physiological conditions, the steady-state platelet count in healthy humans ranges from 150-400 x 10^3^ platelets/µL, with a daily production and clearance of approximately 10^11^ platelets. The hepatic thrombopoietin (TPO) plays a pivotal role as the principal regulator governing the differentiation and maturation of megakaryocytes (MK) in the bone marrow [1]. Diverse mechanisms contribute to platelet clearance, including the hepatic Ashwell-Morell receptor (AMR), which comprises two subunits, ASGPR1 (CLEC4H1, hepatic lectin 1) and ASGPR2 (CLEC4H2, hepatic lectin 2). As platelets age, they undergo desialylation in the bloodstream and are subsequently sequestered in the liver tissue by hepatocytes and Kupffer cells through AMR binding [6, 7], activating Jak2 (Janus kinase 2) and culminating in the removal of senescent platelets [8]. The activation of Jak1 and Jak2 is interleukin-induced, resulting from the binding of interleukin (IL)-6 to the hepatic IL-6 receptor (IL-6R) [9], which induces the phosphorylation of signal transducer and activator of transcription (STAT) 3, facilitating TPO mRNA biosynthesis for the release of TPO into the plasma, thereby concluding the platelet’s life cycle [1, 6].

To elucidate the complex mechanisms governing platelet count regulation following myocardial infarction, we employed a multifaceted approach utilizing several murine models. Specifically, we induced myocardial infarction through ischemia/reperfusion injury (I/R) in wild-type (wt) mice, as well as in mice with thrombocytopenia and GPVI knock-out mice, to investigate the role of STAT signaling in the liver tissue on platelet count regulation. By comparing these models, our study aimed to provide novel insights into the molecular pathways underlying the dynamic changes in platelet counts observed after myocardial infarction. Our findings suggest that the AMR-IL-6R-induced TPO signaling pathway plays a crucial role in regulating platelet turnover and maintaining steady-state platelet counts after MI, thereby ensuring hemostasis.

## 2 METHODS

### 2.1 Animals

All animal experiments complied with the ARRIVE guidelines and were carried out in accordance with the European Parliament guidelines for the use of live animals in scientific studies and the German law for animal welfare. Animal experiments were conducted in accordance with the Declaration of Helsinki and approved by the Ethics Committee of the Ministry of Agriculture, Food and Forestry of the State of North Rhine-Westphalia, Germany. Approval for the experimental protocol was obtained from the Animal Care Committee at Heinrich-Heine-University and the district government of North-Rhine-Westphalia (LANUV; NRW; Permit Numbers AZ 84-02.04.2015.A558; AZ 81-02.04.2019.A270). Only male mice were used in this study.

Mice lacking GPVI (*Gp6*^-/-^) were provided by J. Ware (University of Arkansas for Medical Sciences, Little Rock, AR) and backcrossed to C57BL/6J mice. Heterozygous breeding partners were mated to generate homozygous wild-type control (*Gp6*^*+/+*^) mice and (*Gp6*^*-/-*^) mice. C57BL/6J mice were purchased from Janvier Labs (Le Genest-Saint-Isle, France). Mice were kept in an environmentally controlled room at 22 ± 1°C and a 12 h day-night cycle. The mice were housed in type III Makrolon cages and had *ad libitum* access to food (standard diet) and water. To induce thrombocytopenia, platelet depletion was initiated by injecting a GPIbα antibody (mouse platelet depletion antibody, #R300, polyclonal anti-GPIb alpha, Emfret, Eibelstadt, Germany) or the corresponding IgG control antibody (#C301, polyclonal non-immune rat immunoglobulins (IgG), Emfret, Eibelstadt, Germany) 24 hours before ligating the left anterior descending artery (LAD), and on subsequent days as indicated. According to the data sheet (#R300, Emfret, Eibelstadt, Germany), mice received 2 µg/g of the antibody, dissolved in sterile PBS, to deplete platelets. Platelet depletion, general blood cell count at various time points after I/R were monitored using the automated hematology analyzer Sysmex (Sysmex Corporation, Kobe, Japan).

### 2.2 Experimental model of Acute Myocardial Infarction (AMI) and reperfusion in mice

A closed-chest model of reperfused myocardial infarction was employed to minimize surgical trauma and subsequent inflammatory reactions from the intervention and antibody injection following I/R [10]. Male mice aged 10 to 12 weeks were anesthetized with Ketamin (100 mg/kg body weight, Ketaset®, company: Zoetis, Malakoff, France) and Xylazin (10 mg/kg body weight, Xylazin, company: WDT, Ulft, Netherlands) via a singular intraperitoneal (i.p.) injection before surgery. Euthanasia was carried out through cervical dislocation.

Following successful anesthesia, the left anterior descending artery (LAD) was ligated for 60 minutes to induce MI 3 days post instrumentation. Coronary occlusion was achieved by gently tightening the applied suture until ST-elevation appeared on the electrocardiogram (ECG). Reperfusion was confirmed by the resolution of ST-elevation. Left ventricular function after MI was assessed through echocardiography at different time points post I/R (1 d, 5 d, 21 d) using the Vevo 3100 ultrasound machine (VisualSonics Inc., Bothell, WA, USA). Ejection fraction (%) was measured with corresponding software to confirm reduced LV function in mice.

### 2.3 Measurement of platelet counts

For determining the platelet count whole blood was acquired by puncturing the retrobulbar vein plexus of isofluran anesthetized mice. For anesthesia mice were supplied with 2-3% isoflurane (Isofluran-Piramal, Piramal Critical Care B.V., Voorschoten, The Netherlands) and an oxygen flow rate of 1 L/min. Blood was collected in 300 µL heparin solution (20 U/mL in PBS, Ratiopharm, Ulm, Germany) at different time points after MI. Platelets were counted by using the automated hematology analyzer Sysmex (Sysmex Corporation, Kobe, Japan).

### 2.4 Flow cytometric analysis of platelets

For flow cytometric analysis murine blood was collected in heparin solution (20 U/mL, Ratiopharm, Ulm, Germany) and washed twice by centrifuging 5 min at 650 g with Tyrode‘s buffer (134 mM NaCl, 12 mM NaHCO_3_, 2.9 mM KCl, 0.34 mM Na_2_HPO_4_,20 mM HEPES, 10 mM MgCl_2_, 5 mM glucose, 0.2 mM CaCl_2_, pH 7,35). The blood was resuspended in Tyrode‘s buffer containing 1 mmol/L CaCl_2_. To analyze the ratio between aged and newly formed platelets after myocardial infarction fluorescence based flow cytometry was performed. The number of reticulated platelets was determined by Thiazole Orange staining with retic-count reagent (BD, Franklin Lakes, USA) and GPIbα-antibody (CD42b, Emfret Analytics, Eibelstadt, Germany) to detect newly formed platelets. GPIbα positive cells were gated accordingly to their forward and side scatter profile and were analyzed as described elsewhere [6]. With corresponding software FlowJo (BD, Ashland, USA) the geometric mean of the FSC signal of platelets was acquired additionally. For determining desialylated platelets, platelets were incubated with FITC-labeled lectin (RCA-1, No. FL-1081, Fluorescein (*Ricinus communis*) Agglutinin 1, Vector Laboratories Inc., Burlingame, USA) to detect desialylated platelets. Phosphatidylserine (PS) -exposure of platelets after myocardial infarction was determined by Annexin-binding (Cy™5 AnnexinV, BD Pharmingen, franklin Lakes, USA) and a co-labeling with FITC-conjugated GPIbα-antibody (FITC-conj. rat anti-mouse GPIbα (CD42b), Emfret Analytics, Eibelstadt, Germany) while high Calcium binding puffer (10 mM HEPES, 140 mM NaCl, and 2.5 mM CaCl_2_, pH 7.4) was used.

### 2.5 Immunohistochemistry of spleen sections

At different time points after myocardial infarction spleens were removed, embedded in paraffin and cut into 5 µm sections using an automatic microtome (Microm HM355; Thermo Fisher Scientific, Dreieich, Germany). Paraffin-embedded spleen sections were stained by Hematoxylin/Eosin (H/E) solution (Carl Roth, Karlsruhe, Germany) and the total number of megakaryocytes was counted per visual field.

### 2.6 Quantitative Real-Time polymerase chain reaction (qRT-PCR) from liver tissue

For the analysis of endogenously expressed levels of *Tpo, Asgr1* (*Clec4h1*), *Asgr2* (*Clec4h2*) and *Il-6 receptor* (*Il6ra*) total RNA of the liver before, 6 and 24 hours after myocardial infarction was used. Liver tissue was homogenized in 500 µL cold Trizol by using a tissue homogenisator Precellys (Precellys 24-Dual Homogenisator, Bertin, Frankfurt am Main, Germany) and Precellys Lysing Kit (Precellys, P000918-LYSK0-A, Bertin, Frankfurt am Main, Germany) 30 seconds for 5000 rpm. Total liver RNA was isolated by Trizol/chloroform extraction. Therefore 100 µL Chloroform was added to the liver lysates, followed by centrifugation for 15 min at 18000 g at 4°C. The upper aqueous phase was collected and mixed with 100% ethanol for DNA precipitation. The samples were then purified by using the RNAeasy Mini Kit (Qiagen, Hilden Germany), following the manufacturer‘s protocol. Total RNA concentration and purification was measured with an Eppendorf Bio Photometer® D30. For cDNA generation a total of 200 ng RNA was used with Reverse Transcription Kit (InPromII Reverse transcription System; No, A3800; Promega; Walldorf, Germany). After reverse transcription, quantitative PCR amplification was performed using the following oligonucleotide primers:

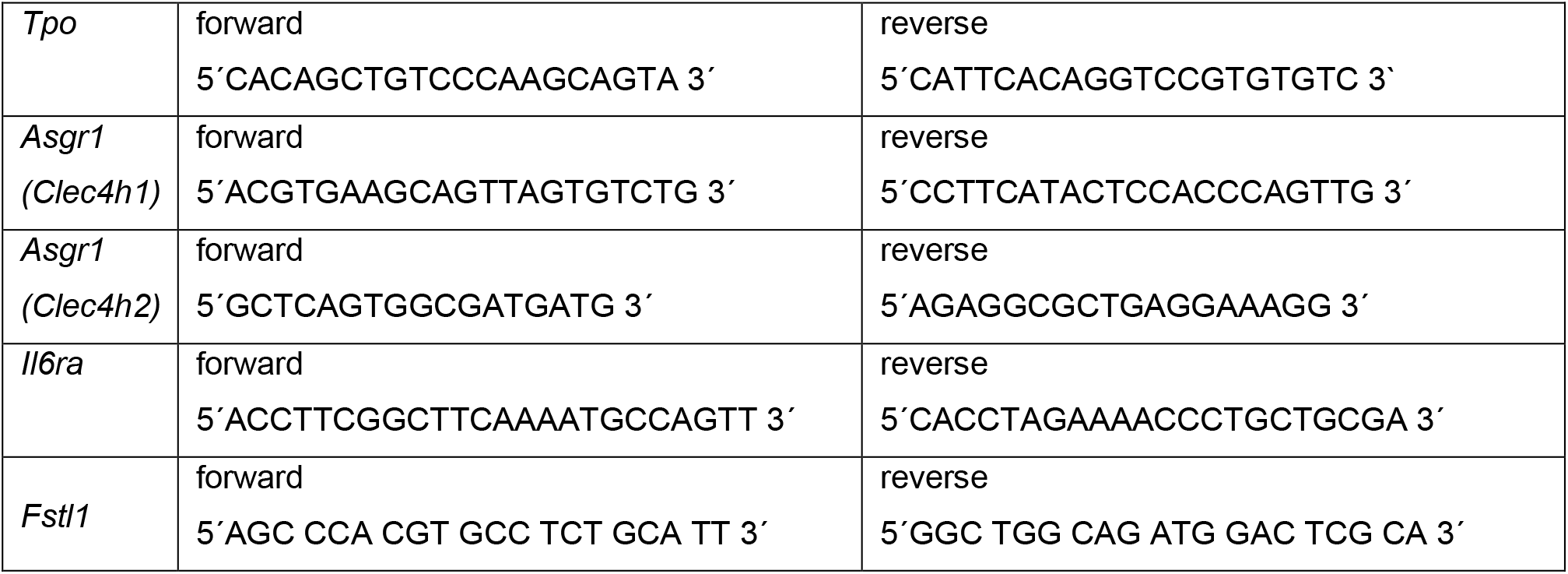

Quantitative Real-time PCR was performed by using Fast Sybr Green Master Mix (Thermo Fischer Scientific, Waltham, USA) following the manufacturer’s protocol. The expression level of the target was normalized to *Follistatin Like 1* (*Fstl1*) RNA expression levels from naive control mice. For analysis of the data the ΔΔCT method was used.

### 2.7 Western Blot (WB) analysis of liver tissue

To determine the expression level of total and phosphorylated form of STAT3 (STAT3 (D3Z26) Rabbit mAb (No. 12640 Cell Signaling Technology, Danvers, MA, USA) P-STAT3 (Y705) Rabbit mAb, No. 9145 Cell Signaling Technology, Danvers, MA, USA), Western Blot analysis of liver lysates before and after myocardial infarction was performed. Liver was homogenized in RIPA buffer (150 mM NaCl, 50 mM Tris-Base, 0.1% SDS, 0.5% sodium deoxycholate, 1% TritonX-100, 1x protease inhibitor complete (Roche, Germany), pH=8.0) by using the tissue homogenisator Precellys (Precellys 24-Dual Homogenisator, Bertin, Frankfurt am Main, Germany) and Precellys Lysing Kit (P000818-LYSK0-A, Bertin Frankfurt, Germany) for 30 seconds and 5000 rpm. After determination of protein concentration, lysates were prepared with a concentration of 50 µg/sample in reducing sample buffer (Laemmli buffer), denaturated for 5 min at 95°C, separated on SDS (sodium dodecyl sulfate)-polyacrylamide gel and transferred onto nitrocellulose blotting membrane (GE Healthcare, Hercules, USA). All antibodies were from Cell Signaling (Cambridge, UK) and usage was followed by manufacturer‘s protocol. For visualization membranes were incubated with peroxidase-conjugated goat-ant rabbit IgGs (GE Healthcare, Chicago, USA) and Immobilon™ Western Chemiluminescent HRP Substrate solution (BioRad, Hercules, USA). GAPDH (14C10, No. 2118, Cell Signaling Technology) served as a loading control.

### 2.8 Enzyme-linked immunosorbent Assay (ELISA)

For quantification of IL-6 and TPO in murine plasma at different time points after myocardial infarction, heparinized blood was centrifuged 10 min for 650 g to retrieve the plasma. The cytokine amount was measured via manufacturer’s protocol (Mouse IL-6 DuoSet ELISA, DY406; Mouse Thrombopoietin DuoSet ELISA, DY488; R&D Systems).

### 2.9 Statistical analysis

All experiments were performed at least three times with n defined as individual animal. Data are presented as arithmetic means ± SEM (Standard Error of Mean) as indicated. Statistical analysis was performed using statistic analyzing software GraphPad Prism 9.5.1 (GraphPad Software, Inc, San Diego, USA). For statistical differences between two groups unpaired student’s *t*-test or unpaired multiple *t* test was performed. For statistical analysis for more than 2 groups one-way or two way ANOVA followed by Tukey’s, Sidak’s or Dunnett’s post hoc multiple comparison test was performed. Statistical tests were performed with P < 0.05 classified as significant. For all figures asterisks or rhombus represent significant differences (*, # p < 0.05; **; ## p < 0.01; ***, ### p < 0.001).

## 3 RESULTS

### 3.1 Acute myocardial infarction leads to an enhanced platelet turnover

C57BL/6J mice underwent acute myocardial infarction (AMI) by ligation of the LAD for 60 min. followed by reperfusion for 24h, 7 and 21 days. A closed-chest model of AMI was employed to minimize surgical trauma and subsequent inflammatory reactions from the intervention post I/R. Ejection fraction as marker for cardiac dysfunction was determined by echocardiography in healthy mice and after 24h and 7 and 21 days post I/R to confirm successful injury of the left ventricle (Figure 1A). Determination of platelet counts in healthy mice and 6 to 24h and 5 to 21 days post I/R revealed a significant reduction only after 6h while normal platelet counts were already achieved after 24h and remained stable until the end of the observation period of 21 days (Figure 1B). The geometric mean of GPIb positive platelets isolated from whole blood of mice at indicated time points showed a significantly enhanced platelet size 6h that turned to be reduced after 24h post I/R (Figure 1C). The number of reticulated platelets was elevated at 6h post I/R compared to healthy controls as determined by thiazolorange staining (Figure 1D). Next, we examined the sialic acid content of circulating platelets post AMI. The fraction of desialylated platelets was examined by platelet lectin binding, such as *Ricinus communis* agglutinin (RCA-1), a lectin that recognizes terminal galactose residues (Figure 1E). 6h after I/R, we detected elevated lectin binding compared to healthy controls, reflecting an enhanced percentage of desialylated platelets early after AMI (Figure 1E). In contrast, no differences were observed at 24h or 5 days post I/R compared to controls. In addition, we determined the number of Annexin-V positive platelets at different time points. As shown in figure 1F, an enhanced number of apoptotic platelets was detected at 24h post I/R. Moreover, early after I/R, we detected pre-activated platelets under non-stimulating conditions (resting) platelets as detected by the binding of JON-A that detects active integrin αIIbβ3 and by elevated P-selectin exposure at the platelet surface at 4h post I/R (Figure 1G-H). However, after 21days, basal platelet activation was significantly reduced compared to naïve controls.

**Figure 1.**
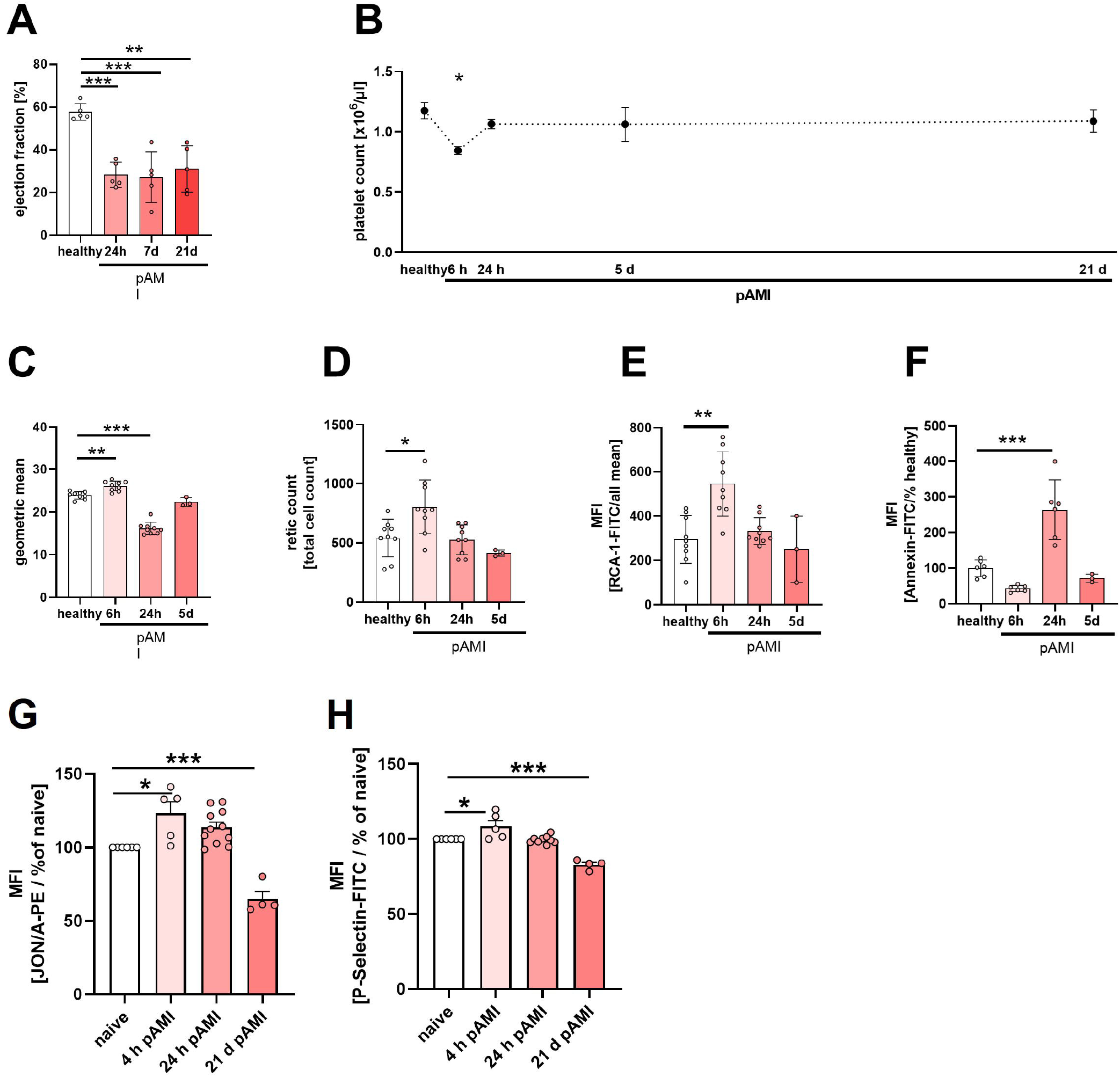
Myocardial infarction leads to enhanced platelet turnover. Mice underwent ischemic/reperfusion (I/R) injury using a closed-chest model. Left ventricular function after I/R injury was assessed through echocardiography. (A) Ejection fraction was determined at indicated time points after I/R to confirm myocardial injury (n=4). (B) Platelet count was measured in whole blood of WT mice before and after I/R injury (n=12 (healthy), 7 (6h), 12 (24h), 7 (5 d), 10 (21d)). (C) GPIb positive platelets in washed whole blood were analyzed with regard to size as determined by the geometric mean of the FSC scatter profile (n=9). (D) The number of reticulated platelets was determined via thiazolorange staining using flow cytometry (n=9). (E) Desialylated platelets were analyzed regarding their *Ricinus communis* agglutinin-1 (RCA) binding profile (n=9). (F) Apoptotic platelets were detected by AnnexinV binding (n=6). (G-H) Activation of platelets at indicated time points was determined by using the antibodies JON/A-PE that binds to active integrin αIIbβ3 (G) and P-selectin as marker for platelet degranulation (H) (n=5-10). Data are presented as means ± SEM. Statistical analyses were done via ordinary one-way ANVOVA with Tukey’s multiple comparison test. * p < 0.05, ** p < 0.01, *** p < 0.001.

### 3.2 Elevated thrombopoietin expression early after ischemia and reperfusion injury

We next analyzed the consequences of elevated numbers of reticulated and desialylated platelets early after AMI and the mechanism how platelet count and function was restored after 24h post I/R. To this end, we first determined TPO plasma levels and detected significantly elevated TPO in the plasma of mice after 24h post I/R compared to healthy controls (Figure 2A). Desialylated platelets bind to the AMR in the liver to induce the phosphorylation of JAK2 and the acute-phase response modulator STAT3. Therefore, we investigated the phosphorylation of STAT3 at different time points post I/R. As shown in figure 2B, we detected significantly elevated activation of STAT3 after 6h post I/R that returns to levels of naïve mice already after 24h. Consequently, no differences in STAT3 phosphorylation between naïve controls and mice with AMI were detected after 24h and 5 and 21 days post I/R (Figure 2B)

Megakaryopoiesis is mainly controlled by the AMR-JAK2-STAT3-TPO signaling pathway in the liver [6, 8]. Therefore, we determined the expression of the AMR subunits Asgr1 and Asgr2 after AMI. We found elevated Asgr1 expression at 6 and 24h after AMI by trend that was significantly reduced thereafter resulting in unaltered gene expression of Asgr1 after 5 and 21 days post I/R compared to healthy controls (Figure 2C). In contrast, expression of the Asgr2 subunit was significantly elevated after 24h post I/R compared to controls.

**Figure 2.**
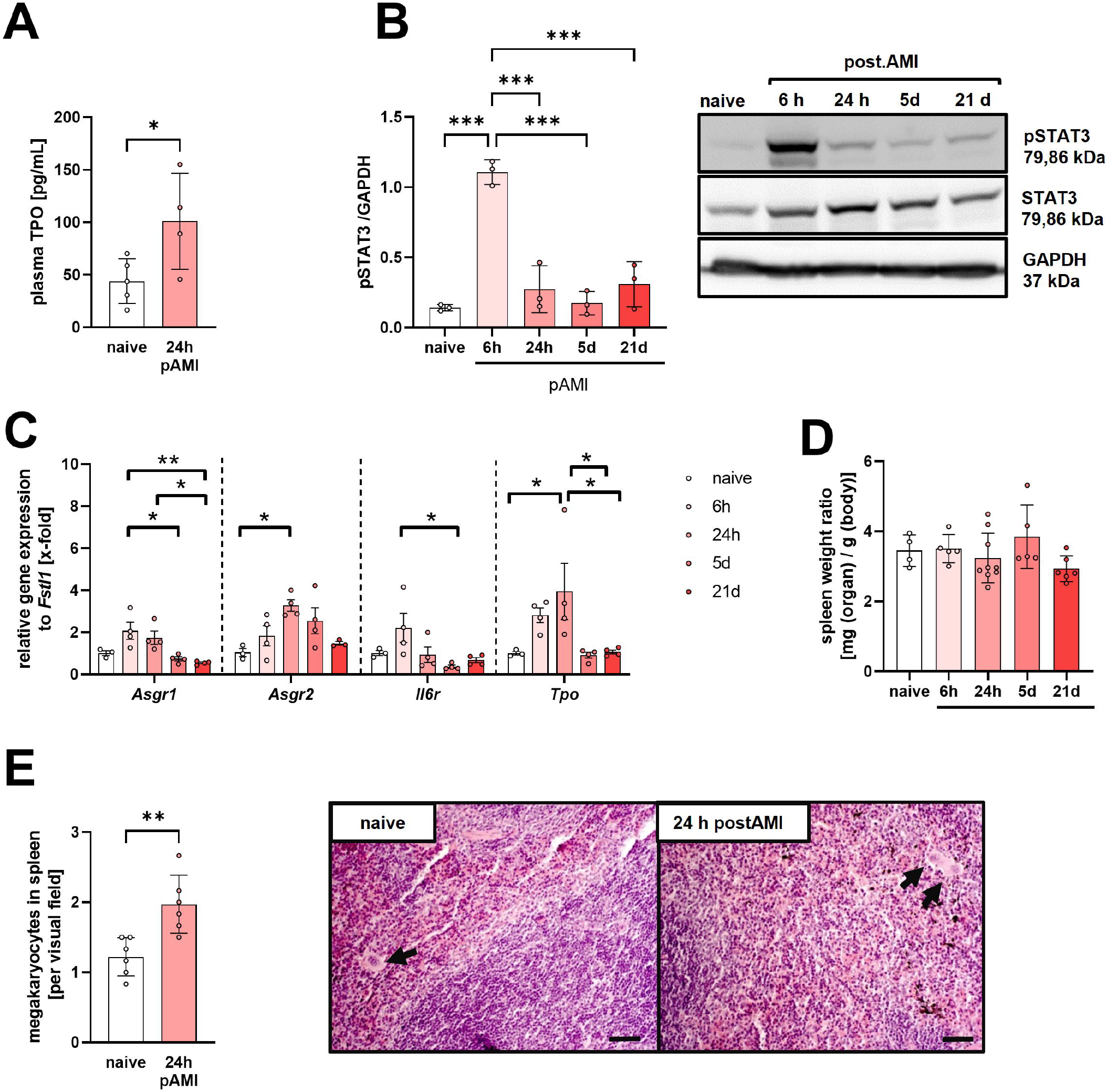
Enhanced expression of TPO early after myocardial infarction. (A) Analysis of plasma thrombopoeitin (TPO) concentrations in WT mice before and 24h after I/R injury (n= 4). (B) Whole liver tissue of WT mice was analyzed for the phosphorylation of STAT3 at indicated time points after I/R injury (n=3). (C) Relative gene expression of different receptors involved in the regulation of *Tpo* expression was analyzed in whole liver tissue of WT mice after I/R injury at indicated time points. (n = 3-4). (D) Relative spleen weight was measured after I/R injury in mice (n= 5-9). (E) Spleen sections were prepared and stained with H&E to count the total number of megakaryocytes per FOV (n=6). Data are presented as means ± SEM. Statistical analyses were done via unpaired student’s t-test (A+E) or ordinary one-way ANVOVA with Tukey’s multiple comparison test (B-D). * p < 0.05, ** p < 0.01, *** p < 0.001. Scale bar =20 µm. H&E = hematoxylin & eosin; FOV = Field of view.

We recently provided evidence for the IL-6 receptor to contribute to JAK2-STAT3-TPO signaling after liver injury by upregulation and crosstalk with the AMR [6]. After AMI, we detected elevated expression of the IL-6 receptor in liver tissue by trend that was significantly reduced compared to late time points post I/R (5 days) (Figure 2C). In line with elevated expression of the AMR1/2 and the IL-6 receptor, and increased phosphorylation of STAT3, we detected increased expression of Tpo in liver tissue after 6 and 24h post I/R that was significantly reduced after 5 and 21 days post AMI (Figure 2C).

Next, we were able to provide first evidence that elevated expression and signaling via the AMR/IL-6-STAT3-TPO pathway affects megakaryopoiesis in spleen because the number of megakaryocytes in spleen was significantly upregulated while spleen weight to body weight ratio was unaltered between naive controls and mice that underwent AMI (Figure 2D-E).

### 3.3 Acute thrombocytopenia triggers megakaryopoiesis after acute myocardial infarction

To provide direct evidence for (desialylated) platelets to be responsible for elevated AMR-IL6-receptor-STAT3 signaling in the liver and the consequences of thrombocytopenia for increased platelet turnover after AMI, we used a thrombocytopenic mouse model. Mice were injected with a platelet-depleting antibody to reduce platelet counts to <1% of controls (Figure 3A). No differences were observed in spleen weight (Figure 3B). However, plasma IL-6 cytokine levels were reduced in thrombocytopenic mice confirming recent observations that platelet play a major role in the increase of IL-6 plasma levels upon acute inflammation [11]. In addition, we determined plasma TPO levels in mice after 6h of I/R. As shown in figure 3D, TPO plasma content was significantly enhanced in thrombocytopenic mice compared to controls (Figure 3D). Although we detected elevated *Tpo* expression in the liver of platelet depleted mice in the naïve state, we did not observe significant differences after 6 and 24h post I/R between the groups (Figure 3E, left panel). However, *Tpo* expression persists at high level in platelet depleted mice after 24h post I/R while it was already decreased in IgG control mice. In contrast, the increase in the expression of the AMR subunit Asgr1 and Asgr2 after 6 h of I/R strongly declines in thrombocytopenic mice to almost basic levels after 24h of I/R while the reduction in Asgr1 and Asgr2 expression was less pronounced in control mice. Thus, a significant difference in the expression of both AMR subunits between platelet depleted and control mice was observed after 24h of I/R (Figure 3E, middle panels). The determination of IL-6R expression in liver tissue revealed a slight decrease between 6 and 24h post I/R in control mice while the decrease was more pronounced in thrombocytopenic mice without reaching statistical significance between the groups at 24h post I/R (Figure 3E, right panel). Next, we analyzed STAT3 phosphorylation in liver tissue. As shown in figure 3F, we detected a significant up-regulation of phosphorylated STAT3 in the liver of IgG control mice, that was less pronounced in platelet depleted mice after 6h of I/R (Figure 3F). No significant differences were observed between groups, neither in naïve mice nor at 24h of I/R, although an enhanced level of phosphorylated STAT3 was observed in naive thrombocytopenic mice compared to naïve controls without reaching statistical significance. However, no significant differences were observed in the number of megakaryocytes in spleen between groups (Figure 3G).

**Figure 3.**
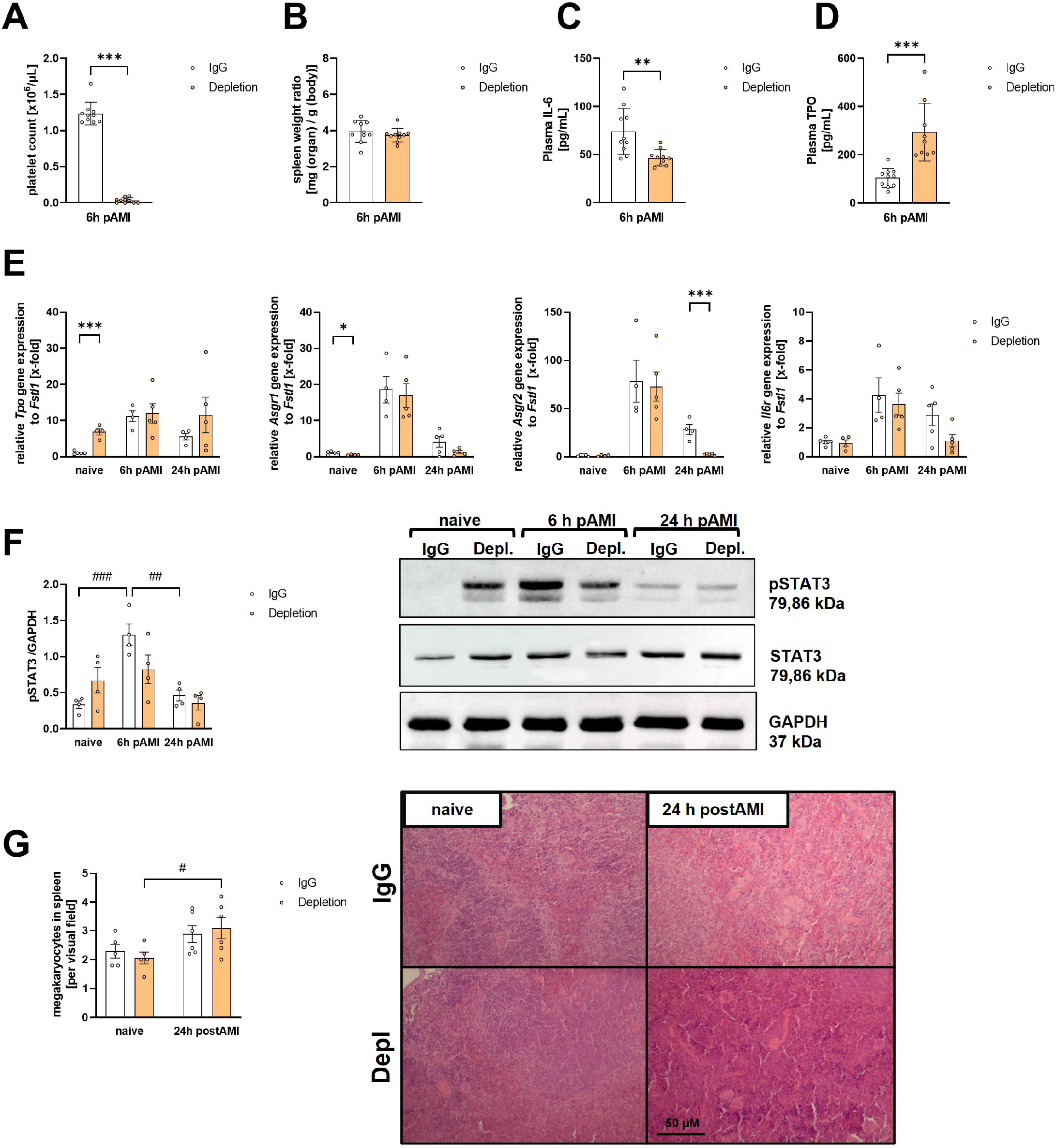
Acute thrombocytopenia triggers megakaryopoiesis after I/R injury. Mice underwent antibody induced platelet depletion 24 hours before LAD ligation. (A) Determination of platelet counts 6h after I/R injury detected in whole blood of mice (n=10). (B) Determination of relative spleen weight ratios 6h after I/R injury in platelet depleted and respective control mice (n=10). Evaluation of IL-6 (C) and TPO (D) plasma levels of platelet depleted mice and respective controls after I/R injury (n=9-10). (E) Relative gene expression of different receptors involved in the regulation of Tpo expression were analyzed in whole liver tissue of platelet depleted mice and respective controls before and after I/R injury (n=4-5). (F) Whole liver tissue was analyzed for STAT3 phosphorylation at indicated time points using platelet depleted mice and respective controls before and after I/R injury (n=4). Quantification of pSTAT3 (left panel) and representative Western blots of pSTAT3, total STAT3 and GAPDH (right panel) are shown. (G) Spleen sections were prepared and stained with H&E to count the total number of megakaryocytes per FOV (n=6). Data are presented as means ± SEM. Statistical analyses were done by unpaired students t-test (A-D), unpaired multiple t test (E) or two-way ANOVA with Tukey’s multiple comparison test (F-G). Asterisks indicate the statistical difference between depletion and the respective controls, while rhombus indicates the differences between different time points before or after MI within one group; *, # p < 0.05, **; ## p < 0.01; ***, ### p < 0.001. H&E = hematoxylin & eosin; FOV = Field of view.

### 3.4 Impact of GPVI on Platelet Count Regulation via Hepatic TPO Signaling Post I/R Injury

Previous studies have identified GPVI as a key regulator of platelet-driven cardiac remodeling following myocardial infarction. Specifically, Inhibition of platelet GPVI binding to collagen by Revacept has been associated with reduced inflammation and improved cardiac function [12], while genetic deletion of GPVI only led to altered scar formation but did not interfere with the acute inflammatory response after AMI [13]. To address the hepatic TPO regulation post I/R, we analyzed the expression of key regulatory genes in *Gp6*^*+/+*^ and *Gp6*^*-/-*^ mice and hepatic STAT3 phosphorylation following myocardial infarction.

To investigate the transcriptional regulation of TPO synthesis by GPVI, we examined the relative expression of *Tpo, Asgr1, Asgr2*, and *Il6r* in liver tissue at different time points post I/R. Both, *Gp6*^*+/+*^ and *Gp6*^*-/-*^ mice exhibited a time-dependent upregulation of *Tpo* and *Asgr1*, while *Asgr2* and *Il6r* followed a similar trend but did not reach statistical significance in *GP6*^*-/-*^ mice compared to WT controls (Figure 4A-D).

**Figure 4.**
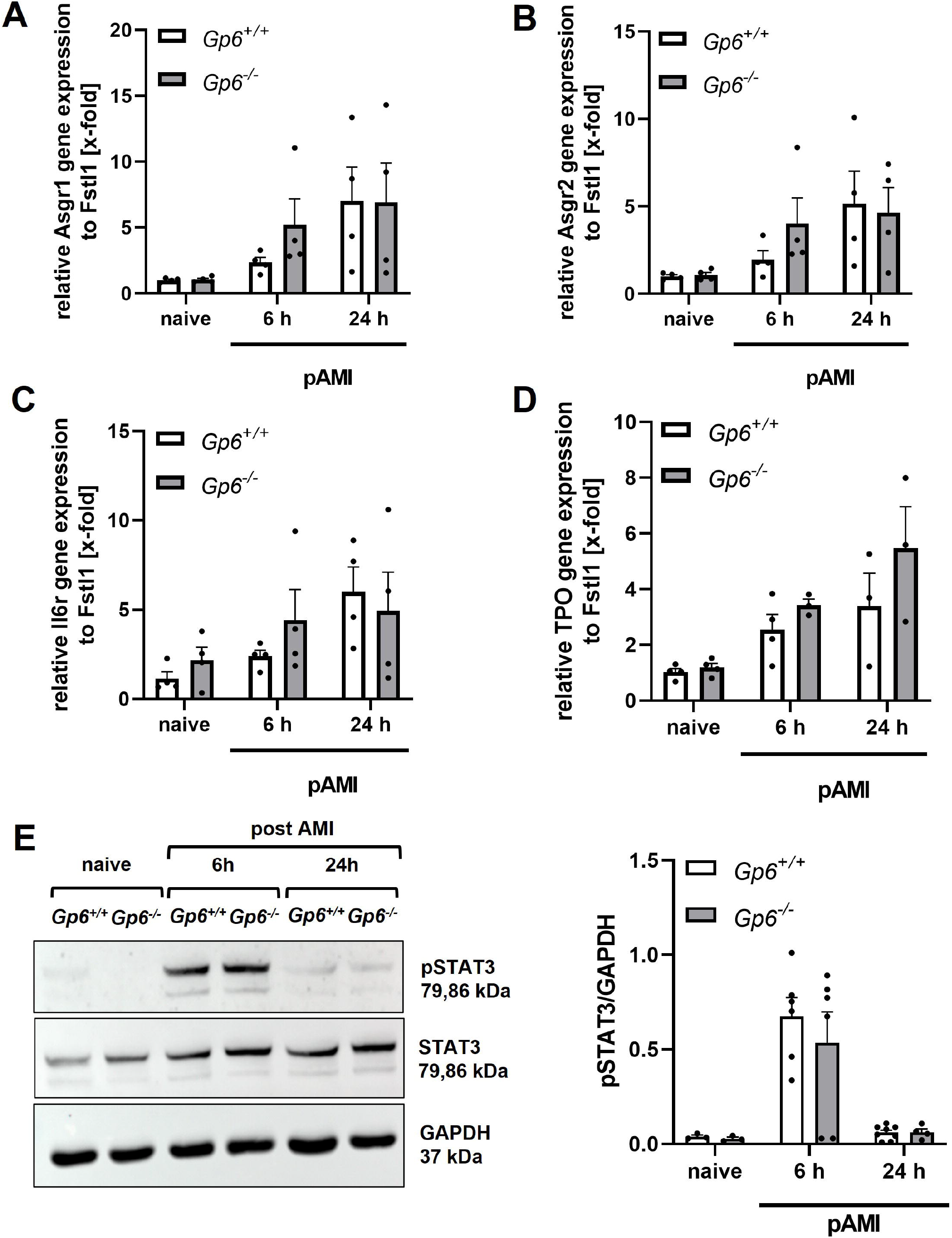
GPVI deficiency has no impact on thrombopoietin production or STAT3 activation in liver tissue at early time points after I/R injury. GPVI deficiency shows no major influence on the platelet turnover after I/R injury. (A) Evaluation of TPO in plasma from *Gp6*^*+/+*^ and *Gp6*^*-/-*^ mice after 24h I/R injury (n=6). (B-E) Relative gene expression of different receptors involved in the regulation of Tpo expression was analyzed in whole liver tissue from *Gp6*^*+/+*^ and *Gp6*^*-/-*^ mice before and after I/R injury (n=3-4). No differences were detected in relative gene expression of Asgr1 (B), Asgr2 (C), IL6r or TPO (E). (F) Phosphorylation of STAT3 was analyzed in whole liver tissue from *Gp6*^*+/+*^ and *Gp6*^*-/-*^ mice at indicated time points before and after I/R injury. Representative Western blots of pSTAT3, total STAT3 and GAPDH (left panel) and quantification of STAT3 phosphorylation (right panel) are shown (n=3-8). Data are presented as means ± SEM. Statistical analyses were done via unpaired student’s t-test (A) or two-way ANOVA with Tukey’s multiple comparison test (B-F).

Next, we assessed hepatic STAT3 phosphorylation, a central regulator of TPO synthesis. In both, *Gp6*^*+/+*^ and *Gp6*^*-/-*^ mice, STAT3 phosphorylation was significantly elevated at 6 hours post I/R, followed by a decline to baseline levels at 24 hours. No significant differences in STAT3 activation were observed between groups, suggesting that GPVI does not contribute to STAT3-mediated TPO induction following myocardial infarction (Figure 4E).

These findings suggest that the absence of GPVI does not significantly impact systemic TPO levels or hepatic STAT3 activation after myocardial infarction.

## 4 DISCUSSION

Platelet hyper-reactivity is a well-documented phenomenon in cardiovascular diseases, including acute myocardial infarction (AMI). Studies have demonstrated that platelets contribute significantly to myocardial reperfusion injury [14, 15]. Enhanced platelet turnover post-AMI leads to an increase in reticulated platelets (RPs), which are linked to worse long-term outcomes [16]. In our study, we observed not only an increase in RPs in WT mice following MI (Fig 1 C+D) in the acute phase after MI, but also a shift in platelet subpopulations with higher frequencies of aged platelets (Fig. 1 E+F). These findings suggest that platelet hemostasis is actively regulated, probably via hepatic regulation of TPO post AMI. To investigate the regulation of TPO after AMI and the role of platelets in this process, we employed two experimental models: (1) antibody-mediated platelet depletion to induce acute thrombocytopenia and (2) genetically modified *GP6*^*-/-*^ mice to assess chronic GPVI deficiency.

The liver plays a key role in platelet clearance also after thrombo-inflammation thereby regulating inflammatory responses including TPO synthesis [17]. Our findings demonstrate that hepatic TPO expression is significantly increased in WT mice after AMI, accompanied by enhanced pSTAT3 signaling. Interestingly, expression of AMR as well as IL6R in liver tissue was enhanced after AMI (Fig. 2B-C). These results align with previous studies indicating that the IL-6/STAT3 pathway mediates heart-liver communication in the setting of myocardial injury [18, 19]. In specific, the authors identified that the liver has a protective role in AMI as regulated by the IL-6/STAT3 signaling pathway. This axis directly inhibits mineralocorticoid receptor expression on hepatocytes, which otherwise exacerbates the outcome after AMI. Given that platelets regulate IL-6 secretion of monocytes and macrophages [20, 21], we observed reduced IL-6 Plasma levels accompanied by reduced STAT3 phosphorylation in thrombocytopenic mice (Fig. 3C). Since platelet desialylation influences STAT3 activation via the AMR [8], we observed enhanced platelet desialylation in line with enhanced pSTAT3 signaling in liver tissue 6h after reperfusion. Therefore, we propose that platelet turnover regulates hepatic signaling pathways, further amplifying thrombo-inflammation, directly via binding of desialylated platelets and immune-modulation of hepatic tissue.

Beyond reduced plasma IL-6 levels, thrombocytopenic mice showed a compensatory upregulation of TPO gene expression and plasma TPO levels (Fig. 3D-E). Interestingly, a compensatory upregulation of TPO after thrombocytopenia is not common. It highly depends on the cause of thrombocytopenia. Immune thrombocytopenic patients show lower TPO levels than expected, while thrombocythemia leads to elevated TPO levels [22, 23]. Thus, challenging the idea of a reciprocal correlation of platelet mass to TPO plasma levels. However, STAT3 signaling remained dysregulated in these mice, suggesting that thrombocytopenia disrupts the dynamic regulation of STAT3 phosphorylation (Fig 3F).

However, a similar phenotype was recently described in a model of murine thrombocytopenia. Sun et al provided evidence that Notch1 signaling regulates TPO production in liver tissue [24]. In their model, *hepatocyte specific (HC) Notch1*^*-/-*^ mice were thrombocytopenic with reduced STAT3 phosphorylation and TPO expression in liver tissue. In contrast to downregulated p-STAT3 in *HC-Notch1*^*-/-*^ mice, we here provided evidence for platelet depletion with only minor effects on the expression of TPO related genes after AMI (Fig. 3E). This suggests that the regulatory mechanism of STAT3 is more complex, arising as a secondary effect of thrombocytopenia following AMI.

In contrast, *GP6*^*-/-*^ mice displayed no significant changes in receptor or TPO regulation compared to WT mice (Fig. 4A-D), with only subtle alterations in IL6R expression early after AMI, likely due to slightly increased baseline IL6R levels in *GP6*^*-/-*^ liver tissue. Additionally, no significant differences in STAT3 activation were observed (Fig. 4E). These findings are consistent with previous reports that GPVI deficiency does not significantly impact acute inflammatory responses following AMI but influences long-term cardiac remodeling such as scar formation [13]. Our data suggests that -while acute platelet depletion triggers a compensatory hepatic TPO response-chronic GPVI deficiency does not exert a similar regulatory effect.

Emerging evidence suggests that the spleen plays a crucial role in platelet dynamics post AMI. It has been shown that splenic platelet release contributes to increased size of circulating platelets and inflammation following AMI [25]. CD41 signals in the spleen are reduced post AMI, indicating an enhanced mobilization of platelets into the circulation. Given our findings of an increased platelet turnover post AMI, it is plausible that splenic platelets contribute to the observed changes in platelet subpopulations and hepatic TPO regulation. Our study also raises questions regarding the potential impact of GPVI-targeted therapies on platelet homeostasis. Antiplatelet therapies, such as Revacept (a GPVI-Fc fusion protein), have been shown to reduce infarct size and improve cardiac function post AMI by preventing platelet-endothelial interactions [12]. However, the consequences of GPVI inhibition on platelet production and hepatic TPO regulation remains unclear. Given the role of GPVI in megakaryocyte function and splenic platelet homeostasis, further studies are needed to determine whether GPVI-targeted therapies influence platelet regeneration in addition to their thrombo-protective effects.

## 5 Conclusions

Our study provides novel insights into the dynamic regulation of the platelet turnover and hepatic TPO signaling following AMI. We demonstrate that acute thrombocytopenia leads to a compensatory increase in hepatic TPO production, accompanied by dysregulated STAT3 signaling, whereas chronic GPVI deficiency does not significantly alter these pathways. Furthermore, our findings suggest that splenic platelet mobilization and systemic thrombo-inflammatory responses may contribute to platelet turnover and hepatic adaptation post AMI. Given the emerging role of GPVI-targeted therapies in cardiovascular disease, future research should explore their impact on platelet homeostasis, hepatic TPO regulation, and long-term cardiac outcomes. Understanding the interplay between platelets, liver signaling, and splenic platelet dynamics could refine therapeutic strategies aimed at mitigating thrombo-inflammation while preserving platelet regenerative capacity post AMI.

## ACKNOWLEDGEMENTS

We thank M. Spelleken and N. Salehzadeh for invaluable technical support.

## AUTHOR CONTRIBUTIONS

F.R. designed the experimental setup, performed flow cytometric analysis, designed figures, and wrote the manuscript; B.T. performed experiments, S.G. operated mice and performed echocardiography to confirm I/R. J.W.F and A.P. contributed with technological tools and provided insights into study design and interpretation of results. M.E. designed the study, supervised the experiments, and wrote the manuscript. All authors read, revised, and approved the manuscript.

## RELATIONSHIP DISCLOSURE

The authors declare that the research was conducted in the absence of any commercial or financial relationships that could be construed as a potential conflict of interest.

## DATA AVAILABILITY

Please contact Margitta Elvers for data sharing.

## REFERENCES

1. Grozovsky, R., et al., Regulating billions of blood platelets: glycans and beyond. Blood, 2015. 126(16): p. 1877–84.

2. Chen, Z.M., et al., Addition of clopidogrel to aspirin in 45,852 patients with acute myocardial infarction: randomised placebo-controlled trial. Lancet, 2005. 366(9497): p. 1607–21.

3. Ndrepepa, G., et al., ST-segment resolution after primary percutaneous coronary intervention in patients with acute ST-segment elevation myocardial infarction. Cardiol J, 2012. 19(1): p. 61–9.

4. Fu, W., et al., Myocardial infarction induces bone marrow megakaryocyte proliferation, maturation and platelet production. Biochem Biophys Res Commun, 2019. 510(3): p. 456–461.

5. Kim, B.S., A Sex-Specific Switch in Platelet Receptor Signaling Following Myocardial Infarction. Preprint, 2019.

6. Reusswig, F., et al., Efficiently Restored Thrombopoietin Production by Ashwell-Morell Receptor and IL-6R Induced Janus Kinase 2/Signal Transducer and Activator of Transcription Signaling Early After Partial Hepatectomy. Hepatology, 2021. 74(1): p. 411–427.

7. Deppermann, C., et al., Macrophage galactose lectin is critical for Kupffer cells to clear aged platelets. J Exp Med, 2020. 217(4).

8. Grozovsky, R., et al., The Ashwell-Morell receptor regulates hepatic thrombopoietin production via JAK2-STAT3 signaling. Nat Med, 2015. 21(1): p. 47–54.

9. Eulenfeld, R., et al., Interleukin-6 signalling: more than Jaks and STATs. Eur J Cell Biol, 2012. 91(6-7): p. 486–95.

10. Reusswig, F., et al., Only Acute but Not Chronic Thrombocytopenia Protects Mice against Left Ventricular Dysfunction after Acute Myocardial Infarction. Cells, 2022. 11(21).

11. Wagenhäuser, M.U., et al., Crosstalk of platelets with macrophages and fibroblasts aggravates inflammation, aortic wall stiffening, and osteopontin release in abdominal aortic aneurysm. Cardiovasc Res, 2024. 120(4): p. 417–432.

12. Schönberger, T., et al., The dimeric platelet collagen receptor GPVI-Fc reduces platelet adhesion to activated endothelium and preserves myocardial function after transient ischemia in mice. Am J Physiol Cell Physiol, 2012. 303(7): p. C757-66.

13. Reusswig, F., et al., Platelets modulate cardiac remodeling via the collagen receptor GPVI after acute myocardial infarction. Front Immunol, 2023. 14: p. 1275788.

14. Schultheiss, H.P., et al., Large platelets continue to circulate in an activated state after myocardial infarction. Eur J Clin Invest, 1994. 24(4): p. 243–7.

15. Xu, Y., et al., Activated platelets contribute importantly to myocardial reperfusion injury. Am J Physiol Heart Circ Physiol, 2006. 290(2): p. H692–9.

16. Bongiovanni, D., et al., Role of Reticulated Platelets in Cardiovascular Disease. Arterioscler Thromb Vasc Biol, 2022. 42(5): p. 527–539.

17. An, O. and C. Deppermann, Platelet lifespan and mechanisms for clearance. Curr Opin Hematol, 2024. 31(1): p. 6–15.

18. Kuyama, N., et al., Mineralocorticoid Receptor Blocker Prevents Mineralocorticoid Receptor-Mediated Inflammation by Modulating Transcriptional Activity of Mineralocorticoid Receptor-p65-Signal Transducer and Activator of Transcription 3 Complex. J Am Heart Assoc, 2024. 13(18): p. e030941.

19. Sun, J.Y., et al., An IL-6/STAT3/MR/FGF21 axis mediates heart-liver cross-talk after myocardial infarction. Sci Adv, 2023. 9(14): p. eade4110.

20. Hawwari, I., et al., Platelet transcription factors license the pro-inflammatory cytokine response of human monocytes. EMBO Mol Med, 2024. 16(8): p. 1901–1929.

21. López, M.L., et al., Encapsulated platelets modulate kupffer cell activation and reduce oxidative stress in a model of acute liver failure. Liver Transpl, 2016. 22(11): p. 1562–1572.

22. Griesshammer, M., et al., High levels of thrombopoietin in sera of patients with essential thrombocythemia: cause or consequence of abnormal platelet production? Ann Hematol, 1998. 77(5): p. 211–5.

23. Kosugi, S., et al., Circulating thrombopoietin level in chronic immune thrombocytopenic purpura. Br J Haematol, 1996. 93(3): p. 704–6.

24. Sun, Y., et al., Notch1 regulates hepatic thrombopoietin production. Blood, 2024. 143(26): p. 2778–2790.

25. Gao, X.M., et al., Splenic release of platelets contributes to increased circulating platelet size and inflammation after myocardial infarction. Clin Sci (Lond), 2016. 130(13): p. 1089–104.

