## Supplemental data for "The heart-liver axis controls platelet turnover by hepatic STAT3 phosphorylation and TPO regulation after acute myocardial infarction"

### Supplementary Material

#### 1 Supplementary Figures and Tables

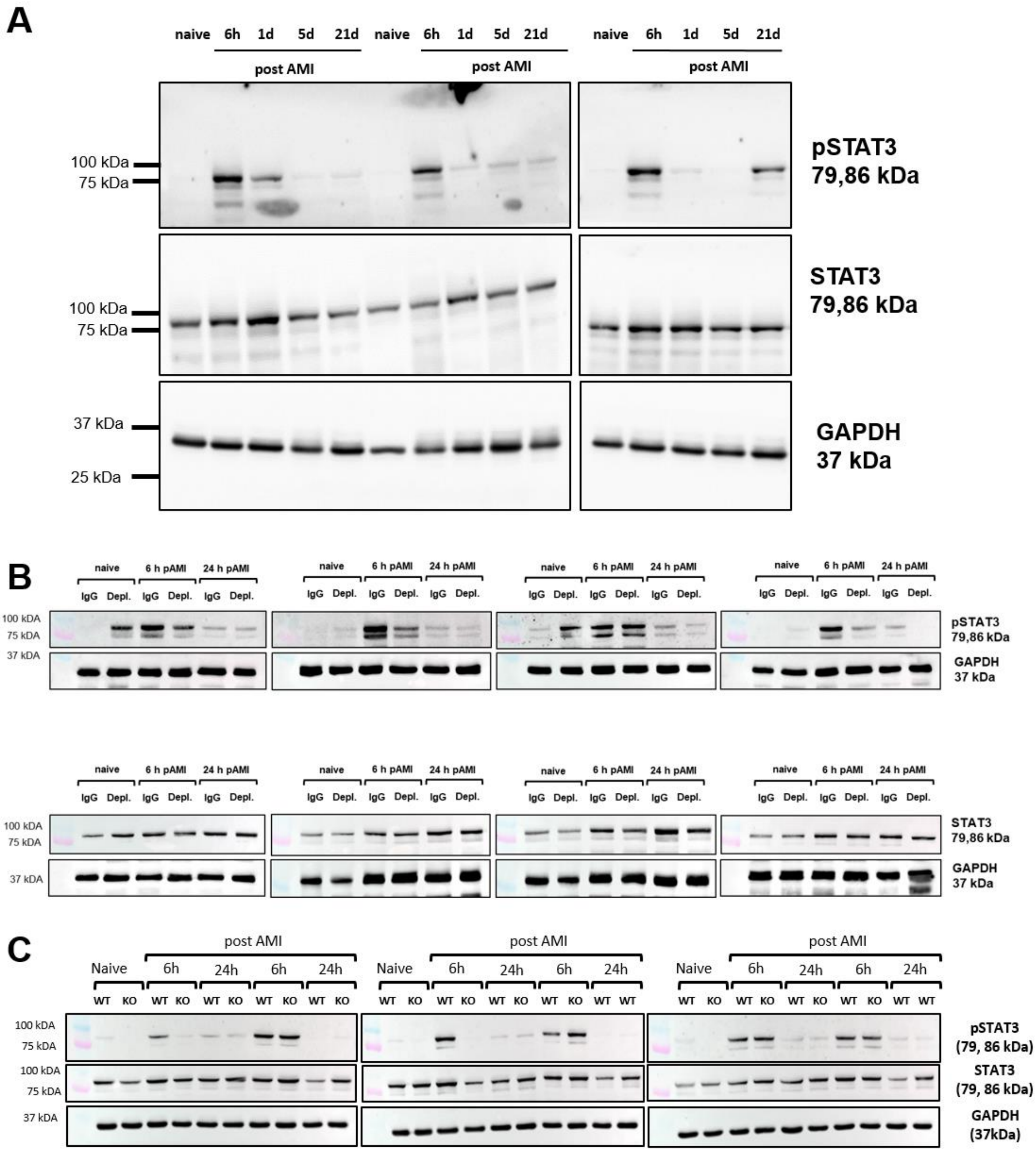


**Supplemental Figure 1** Analysis of STAT3 phosphorylation in whole liver tissue after I/R. Naïve (control) mice and mice that underwent I/R injury were analysed for the phosphorylation of STAT3 and for total STAT3. GAPDH was used as loading control. Analysis of (A) WT mice (n= 3), (B) Platelet depleted and respective IgG control mice (n=4) and (C) $Gp6^{+/+}$ and $Gp6^{-/-}$ mice (n=3).
